# Recognition of an appropriate mating partner using cuticular hydrocarbons in a species with an extreme intra-sexual dimorphism

**DOI:** 10.1101/2024.11.13.623210

**Authors:** Victoria Moris, Aline Wirtgen, Oliver Niehuis, Thomas Schmitt

**Affiliations:** Department of Evolutionary Biology and Ecology, Institute of Biology I (Zoology), University of Freiburg, 79104 Freiburg, Germany; Department of Animal Ecology and Tropical Biology, Biocentre, University of Würzburg, Am Hubland, 97074 Würzburg, Germany

**Keywords:** sexual attractiveness, cuticular hydrocarbons, methyl-branched alkanes, anti-aphrodisiacs

## Abstract

Attracting a mate at the right time is critical for many species that reproduce sexually. In insects, short-range communication between potential mates is often mediated by cuticular hydrocarbons (CHCs), which cover most of the insect cuticle. Although the CHC profiles of many insects have been studied, we know little about what aspects of a CHC profile cause changes in mate attractiveness over the lifetime of an individual. We addressed this question by studying the mason wasp *Odynerus spinipes*, whose females exhibit age-related quantitative changes in their CHC profile composition. First, we created an ethogram of the male mating behavior. We knew from preliminary investigations that males do not attempt to mate with recently eclosed adult females. By coating wasp dummies with different CHC extracts, we were able to show that the CHC profiles of 3-day-old females are indeed more attractive to males than those of 0-day-old females. The increased attractiveness of 3-day-old females compared to younger females was negatively correlated with the relative abundance of methyl-branched alkanes in the CHC profile of the females. These results, and the fact that the CHC profile of males is characterized by a high relative abundance of methyl-branched alkanes throughout the adult wasp lifetime, suggest that in *O. spinipes*, methyl-branched alkanes may act as anti-aphrodisiacs that reduce the harassment of females by males before the females are ready to mate.

## Introduction

Chemical signaling has often been shown to be critical for successful reproduction in various animal species (Wyatt 2014; Blomquist and Ginzel 2021). In particular, insects are known to rely heavily on chemical communication for mate finding and recognition (Wyatt 2014). However, the mechanisms by which specific information, such as sexual attractiveness, is encoded in chemical cues remain poorly understood (Allison and Cardé 2016; Stökl and Steiger 2017).

Cuticular hydrocarbons (CHCs), which cover most of the insect cuticle and protect against desiccation, play an important role in insect sexual communication (Dietemann et al. 2003; Steiger et al. 2011). In most insect species, each sex is characterized by a specific set of hydrocarbons of a certain length (typically 19 to 52 carbon atoms), which may contain double bonds and methyl branches. The pattern of abundance of different hydrocarbons on the cuticle is called the CHC profile. CHCs are known to be used by insects as contact pheromones (Ayasse et al. 2001; Paxton 2005; Ferveur 2005; Kroiss et al. 2006; Niehuis et al. 2013; Wyatt 2014). Qualitative and/or quantitative differences between the CHC profiles of different species and between the sexes of a given species allow individuals to identify conspecific mates (Thomas and Simmons 2008). However, the presence and abundance of specific components in a CHC profile can change over the lifetime of an individual (e.g., (Simmons et al. 2003; Hugo et al. 2006; Ichinose and Lenoir 2009; Nunes et al. 2009; Kuo et al. 2012; Polidori et al. 2017; Bien et al. 2019)). These changes can be indicative of mating status and other traits and qualities of an individual (Paulmier et al. 1999; Ayasse et al. 1999; Schiestl and Ayasse 2000; Simmons et al. 2003; Marcillac and Ferveur 2004; Mant et al. 2005; Grillet et al. 2006; Steiger et al. 2007; Jennings et al. 2014). This, in turn, can influence the attractiveness of the individual (Kuo et al. 2012) and thus whether or not its CHC profile induces a courtship behavior in a potential mate (Finck et al. 2016; Würf et al. 2020).

The mason wasp *Odynerus spinipes* (Eumininae: Vespidae) has proven to be a promising species for elucidating the molecular basis and evolution of CHC profile differences (Moris et al. 2023). Females of this species are known to be capable of expressing one of two CHC profiles, known as chemotypes, which they maintain throughout their adult lives (Wurdack et al. 2015; Moris et al. 2021). The two chemotypes differ qualitatively in 77 CHCs, mainly in alkenes having their double bonds at uneven positions for chemotype 1 and at even positions for chemotype 2 (Wurdack et al. 2015; Moris et al. 2021). In contrast, *O. spinipes* males express only one CHC profile that is very similar to one of the two chemotypes expressed by their females (Wurdack et al. 2015). The composition of the two chemotypes expressed by *O. spinipes* females changes with female age (Moris et al. 2023), but the compositional changes and their functional role have not yet been investigated in detail. Males face the problem of recognizing females with both chemotypes as appropriate mating partner. This could be caused by two sets of compounds, both of which are attractive to males, or by a sex pheromone encoded in the CHC profile of both chemotypes. We hypothesize that females will show a sex pheromone during mating period in both chemotypes that is different from the male CHC profile.

In this study, we report our results from investigating age-related changes in the CHC profiles of O. *spinipes* females and assessing the extent to which these changes alter female attractiveness to conspecific males. Specifically, we (1) recorded the mating behavior of males and females to create an ethogram of the male courtship behavior. Since we know from a preliminary investigation that males do not attempt to mate with recently eclosed adult females, we (2) conducted male O. *spinipes* mating experiments using female O. *spinipes* dummies coated with CHC profiles of females of different ages. Specifically, we compared the attractiveness of CHC profiles of recently eclosed females (0 days old) with those of females that were 3 days old. Finally, given the different response of males to dummies coated with CHC profiles of females of different ages, we analyzed (3) in what aspects the CHC profiles of *O. spinipes* females (both chemotypes) and males (for the sake of completeness) change during the adult life of the wasps, sampling wasps 0, 3, 9 and 14 days after their eclosion.

## Results

### Male mating behavior

We recorded the behavior of nine *O. spinipes* male-female pairs that had eclosed 4–5 days earlier (Fig. 1A; Table 1). We found that the male mating behavior (= B) is highly stereotyped and can be summarized as follows (Fig. 1B–E): when the male physically meets the female, it (B1) makes antennal contact with the female. It then (B2) mounts the female and wraps its antennae around the female’s antennae. Once the female is mounted, the male (B3) extends its aedeagus and rubs it against the female’s metasoma, causing the female to open its genital opening. The male (B4) inserts its genital capsule into the female’s genital opening. After mating with the female, and even after unsuccessful mating attempts, the male (B5) lifts its metasoma and extrudes their genital capsule.

**Fig. 1:**
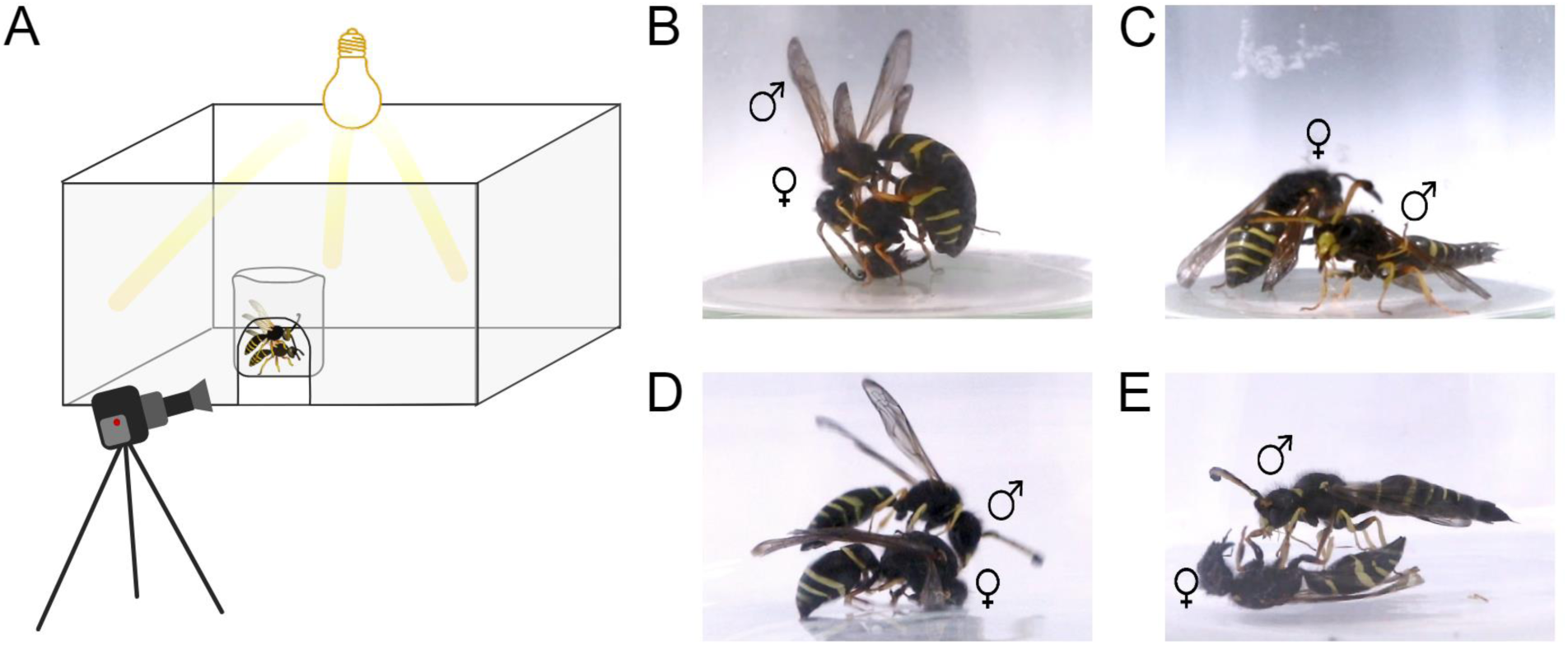
A) Experimental setup for recording the mating behavior of individual *Odynerus spinipes* males when providing a living female or a female dummy coated with the CHCs of females of a specific age class. B) *O. spinipes* male mounts a living female and loops the tips of its antennae around those of the female; C) *O. spinipes* male extruding its genital capsule after having copulated with the female; D) *O. spinipes* male mounts a dummy female; E) *O. spinipes* male extruding its genital capsule after having copulated with a dummy female.

**Table 1:** Occurrences of individual *Odynerus spinipes* male mating behaviors (B1–5) described in detail in the results text. (B1) Male making antennal contact with the female, (B2) male mounting the female and wraps its antennae around the female’s antennae, (B3) male on top of the female extending its aedeagus and rubs it against the female’s metasoma, (B4) male inserting its genital capsule into the female’s genital opening, (B5) male lifting its metasoma and extruding its genital capsule after mating or unsuccessful mating attempts. d = day(s).

| Male | Age | Female | Age | B1 | B2 | B3 | B4 | B5 |
| --- | --- | --- | --- | --- | --- | --- | --- | --- |
| #12922 | 5 d | #12921 | 5 d | + | + | + | + | + |
| #12924 | 5 d | #12923 | 5 d | + | + | + | – | – |
| #12926 | 4 d | #12913 | 4 d | + | + | + | – | + |
| #12928 | 4 d | #12915 | 4 d | + | + | + | + | + |
| #12932 | 4 d | #12919 | 4 d | + | + | + | + | + |
| #12930 | 4 d | #12917 | 4 d | + | + | + | + | + |
| #12936 | 5 d | #12927 | 4 d | + | + | + | + | + |
| #12934 | 5 d | #12925 | 4 d | + | + | + | + | + |
| #12938 | 5 d | #12933 | 4 d | + | + | + | + | + |

### Effect of female CHCs on male attraction

Since we had observed in preliminary studies (Fig. S1) that males did not seem to mate with recently eclosed females, we performed experiments with female dummies whose CHC profile was varied. Female wasp dummies were coated with CHC extracts from *O. spinipes* females of two different age classes (0 days, 3 days). When six males were presented sequentially with two female dummies, each with a different age class CHC coating (order randomized) but the same chemotype, we found that males were significantly more attracted to female dummies coated with CHC extracts from 3-day-old females than to those coated with CHC extracts from 0-day-old females (Fig. 2; paired t-test: p-value = 0.039; Table S1).

**Fig. 2:**
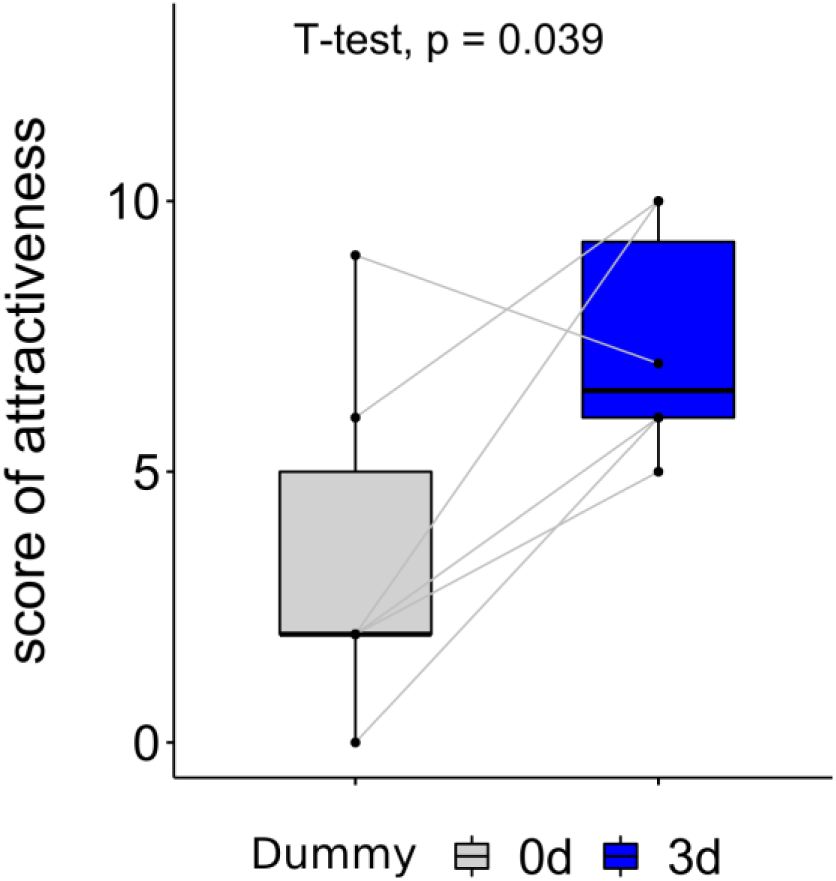
Attractiveness scores of female dummies coated with cuticular hydrocarbons extracted from 0-day-(gray box plot) and 3-day-old (blue box plot) females to six males. Gray lines indicate each pair of data. See Materials and methods for how attractiveness scores were calculated.

### CHC profile differences between 0- and 3-day-old females

We compared the composition of CHC profiles from *O. spinipes* wasps of different age classes (i.e., 0, 1–2, 3–4, 8, 14 days after eclosion), considering both female chemotypes and also the male sex. An NMDS plot of the Bray-Curtis distances of the CHC profiles (Fig. 3) showed that the samples of the two female chemotypes form two clearly distinct groups, with the CHC profiles of the males nested within the group of CHC profiles of the females with chemotype 1. Within each of the two groups, we found that the CHC profiles of female samples from a given age class are clustered and that samples from different age classes are located in different regions of the chemospace. Interestingly, the chemospace of young males largely overlaps with the chemospace of young females with chemotype 1.

**Fig. 3:**
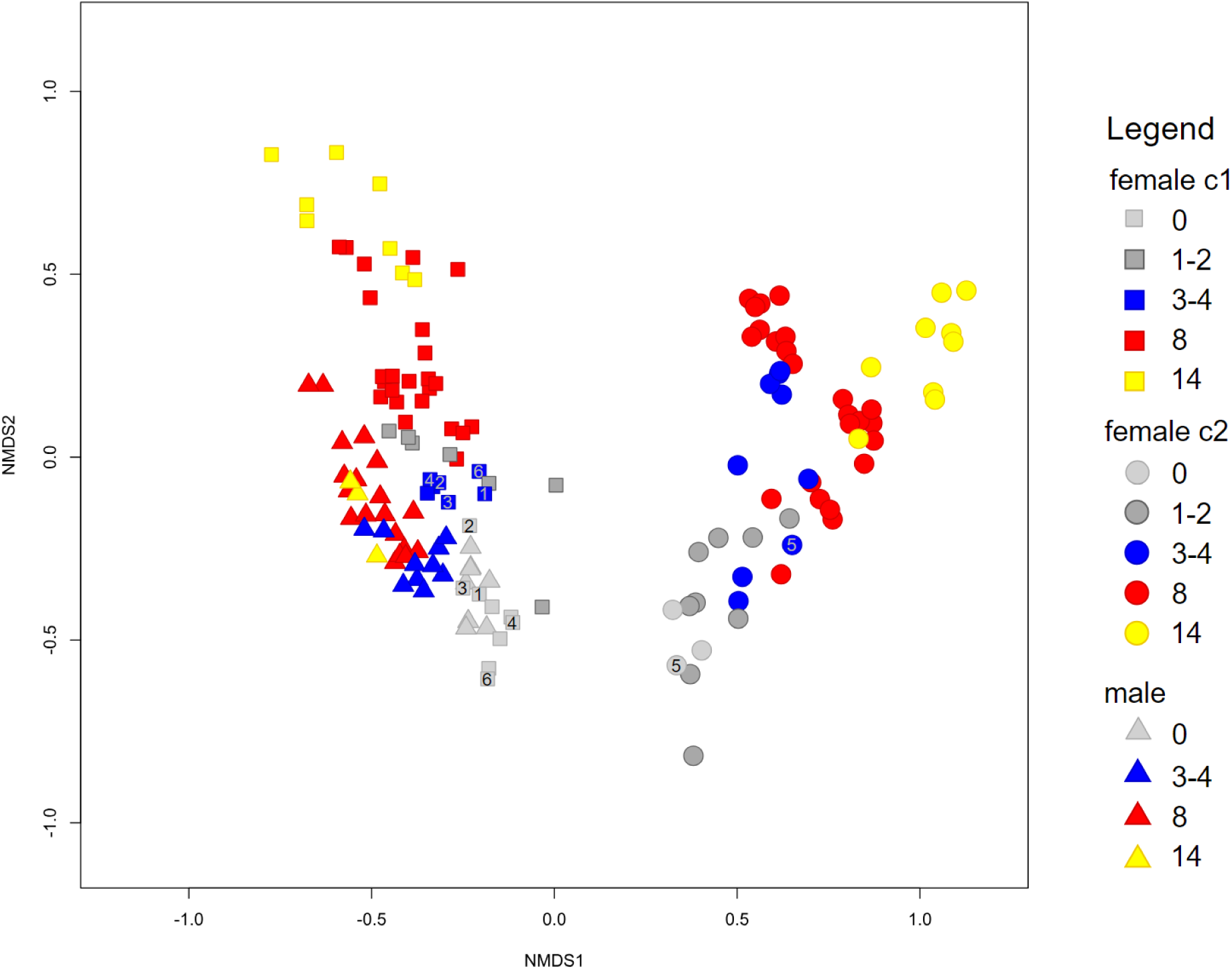
Non-metric multidimensional scaling (NMDS) ordination of Bray-Curtis distances between cuticular hydrocarbon (CHC) profiles of *O. spinipes* males and females. Each data point represents a CHC profile of an *O. spinipes* sample at a different time point: day 0 (gray), day 1–2 (dark gray), day 3–4 (blue), day 7–8 (red), and day 14–15 (yellow). CHC profiles of males (N = 40: N_0d_ =8, N_3d_ = 10 N_8d_ = 19, N_14d_ =3) are indicated by triangles, those of females of chemotype 1 (= c1; N = 55: N_0d_ = 9, N_1d_= 7, N_3d_ = 7, N_8d_ = 24, N_14d_ = 8) by squares and those of chemotype 2 (= c1) females (N = 54: N_0d_ = 3, N_1d_ = 9, N_3d_ = 9, N_8d_ = 24, N_14d_ = 9) by circles. Samples used to cover dummy females are indicated by the ID of the male that had to choose between them (see Table S1).

We used PCA to shed light on the identity of the CHCs driving the differences in CHC profiles between females of the two age classes of particular interest to us (i.e., 0 and 3 days after eclosion) of each chemotype. The first principal component explained a substantial amount of the total variation in CHC profiles (chemotype 1: 42%, chemotype 2: 48%) and clearly separated the CHC profiles of 0-day-old wasps from those of 3-day-old wasps in both analyses (Fig. 4). Analyzing the contributions of each compound to the variability in dimension 1, we found that certain compounds primarily explain the differences in CHC profiles between 0-day- and 3-day-old females, common across both chemotypes but absent in males. These compounds are methyl-branched alkanes: 11-;9-MeC23, x,y-diMeC24, 13-;11-;9-MeC25, and 7-MeC27 (highlighted in red in Fig. 4). Other methyl-branched alkanes, common across both chemotypes and males, include 11,15-diMeC25, 5-MeC25, 11-;9-MeC29, and 11-MeC31 (in green in Fig. 4). Additionally, alkenes contribute to the differences between 0-day- and 3-day-old female CHC profiles, though they vary between chemotypes by double bond position—uneven carbon numbers in chemotype 1 and even carbon numbers in chemotype 2. Some of these alkenes are also present in males (Fig. 4).

**Fig. 4:**
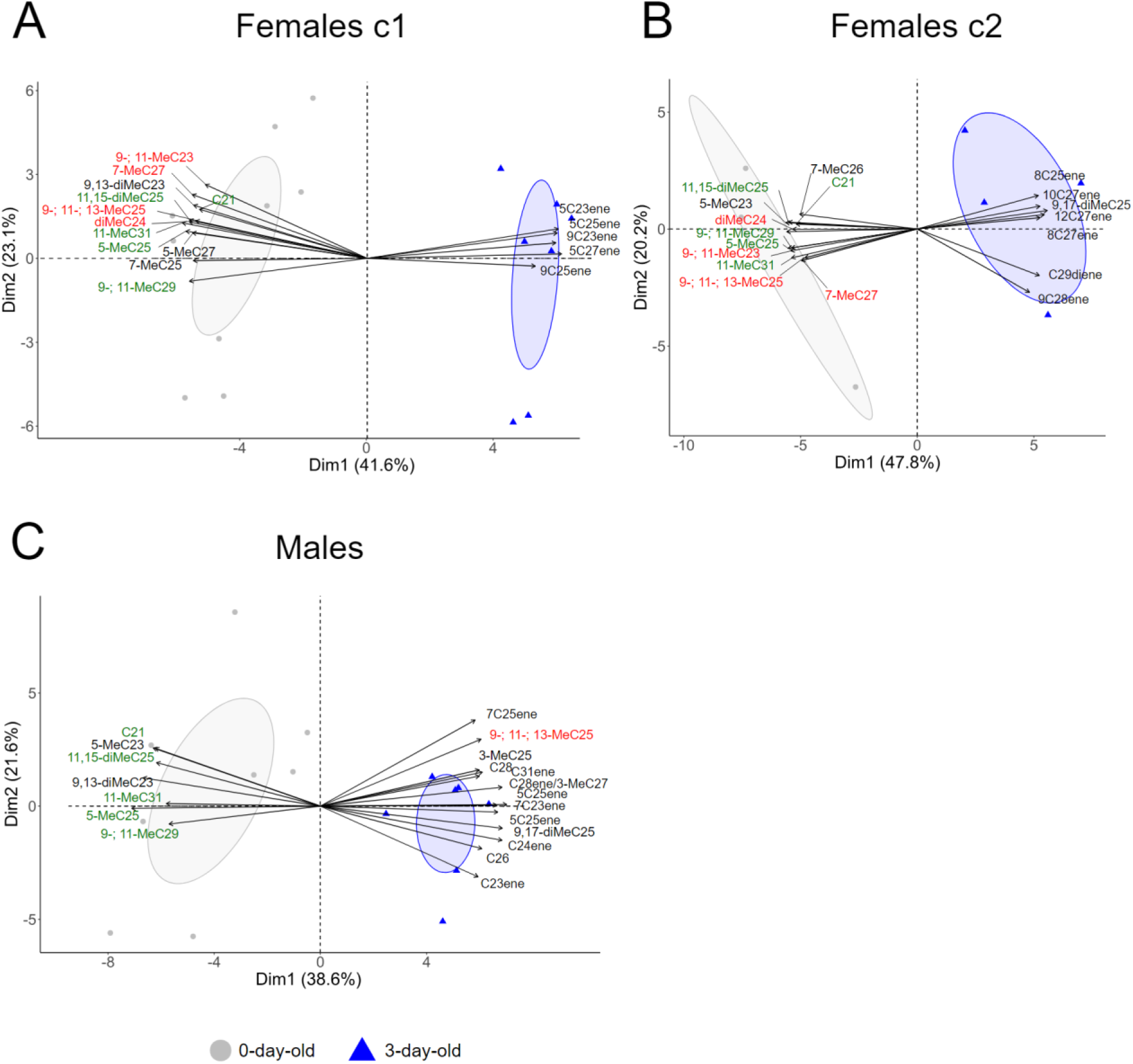
Principal component analysis (PCA) of cuticular hydrocarbon (CHC) profiles of *O. spinipes* females (chemotype 1 [= c1]: left; chemotype 2 [= c2]: middle) and of males, with the eigenvectors of those CHCs whose relative abundances are most correlated along the first principal component. Each marker represents a CHC profile of a wasp of a different age class (grey: day of eclosion; blue: 3 days after eclosion). CHCs that explain the differences between 0-day- and 3-day-old CHC profiles found in common in the two chemotypes but not in male CHC profile are written in red; CHCs that explain the differences between 0-day- and 3-day-old CHC profiles found in common in the two chemotypes and in male CHC profile are written in green.

Given the PCA results, we examined at the overall differences in the abundance of methyl-branched alkanes in the CHC profiles of 0-day- and 3-day-old wasps. Among the two age classes, we found that methyl-branched alkanes differed the most: more than 70% decrease in abundance in the profiles from day 0 to day 3 (chemotype 1: day 0, mean 12.3%, N = 9, day 3, mean 3.7%, N = 7; chemotype 2: day 0, mean 11.6%, N = 3, day 3, mean 3.0%, N = 9). For comparison, males maintained a relatively high abundance of methyl-branched alkanes in their CHC profile over the same time period (day 0, 14.3%, N = 8, day 3, 12.3%, N = 10; Table S2). Indeed, the total abundance of methyl-branched alkanes was significantly reduced for females of chemotype 1 (Wilcoxon test, Bonferroni adjusted p = 0.0009) and for females of chemotype 2 (Wilcoxon test, Bonferroni adjusted p = 0.045) 3 days after eclosion, while it was not significantly different in 0-day- and 3-day old males (Fig. 5, Table S3).

**Fig. 5:**
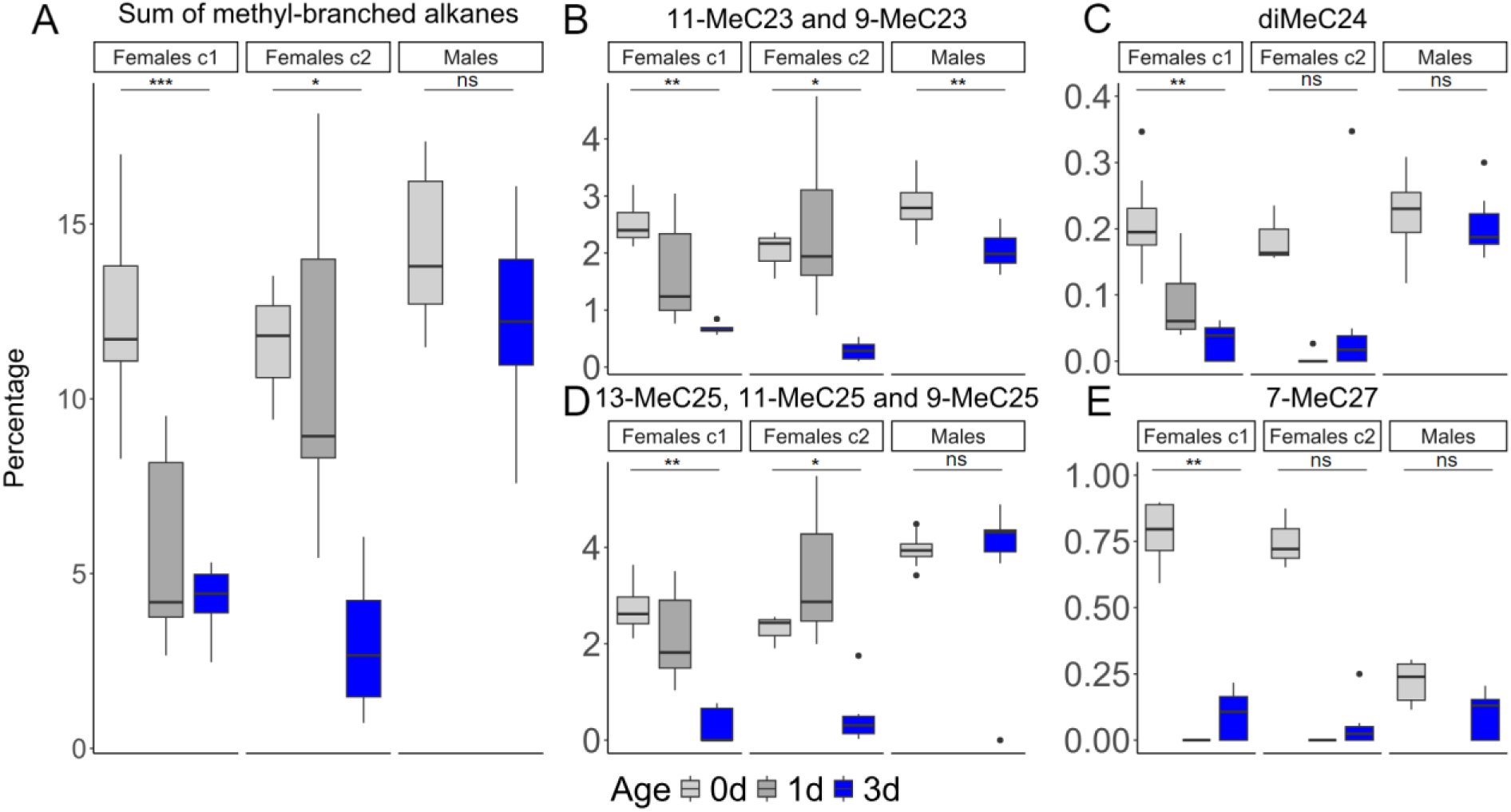
Relative abundance of methyl-branched alkanes in *O. spinipes* males and females (c1 = chemotype 1; c2 = chemotype 2) on the day of eclosion (gray) and 3 days after eclosion (blue). The ordinate shows the relative abundance (in percent) of all methyl-branched alkanes (A), of 11-MeC23 and 9-MeC23 (B), of diMeC24 (C), of 13-MeC23, 11-MeC23 and 9-MeC23 (D), 7-MeC27 (E). Asterisks indicate level of statistical significance of the differences between the two age classes: p <= 0.05, *: p <= 0.01, **: p <= 0.001, ***: p <= 0.0001 (individual p-values and adjusted p-values are provided in Table S3).

Examining the relative abundance of methyl-branched alkanes identified in PCA analyses of females of chemotypes 1 and 2, but absent in males, we observed two distinct groups of compounds that significantly decreased in females of both chemotypes three days after eclosion (Fig. 5, Table S3): 11-; 9-MeC23, and 13-; 11-; 9-MeC25. In males, these compounds maintained a high relative abundance, although the first compound also showed a slight decrease—not as pronounced as in females. These methyl-branched alkanes constitute a major proportion of the overall profiles in both males and females. We also observed a significant decrease in x,y-diMeC24 levels in 3-day-old females of chemotype 1 compared to newly eclosed females, but not in chemotype 2 females, possibly due to a limited sample size (N = 3) for the latter group. Additionally, 7-MeC27 levels significantly declined in 3-day-old females of chemotype 1 compared to their levels at eclosion, with this compound being less abundant in males than in newly eclosed females. Among other highly abundant methyl-branched alkanes (Fig. S2), we found that 3-MeC25 and 3-MeC27 were unique in showing an increase in males, while the remaining compounds declined in both males and females.

## Discussion

We examined age-related changes in the CHC profiles of O. *spinipes* females and assessed the extent to which these changes alter female attractiveness to conspecific males. *O. spinipes* is a particularly interesting species in this regard, as its females can express one of two chemotypes that differ in 77 compounds (Wurdack et al. 2015). If females of the two chemotypes show a similar age-related change in attractiveness to conspecific males, commonalities in their CHC profile changes could point to the causal compound(s).

Using initially odorless dead wasps coated with CHCs extracted from females of different ages and chemotypes, we found that CHC extracts from 3-day-old females of both chemotypes were statistically significantly more attractive to males than those from recently eclosed females. The main difference in CHC composition between recently eclosed and 3-day-old females is the relative abundance of certain methyl-branched hydrocarbons and alkenes. Specifically, methyl-branched alkanes present in the CHC profiles of recently eclosed females are almost absent in females 3 days old or older. The profiles of these older females are also characterized by a relatively high abundance of unsaturated hydrocarbons. The low attractiveness of recently eclosed females may result from a low relative abundance of unsaturated hydrocarbons on their cuticle, a high relative abundance of methyl-branched hydrocarbons, or a combination of both. Although our data do not allow us to test these two hypotheses directly, several observations in this study support the second hypothesis, suggesting that methyl-branched hydrocarbons may act as anti-aphrodisiacs. First, the CHC profiles of males are characterized by a high relative abundance of methyl-branched hydrocarbons, which, unlike in females, does not significantly decrease as males age. Second, although this study did not directly test this, our year-long research on *O. spinipes* found no instances of males attempting to mate with other males. This suggests that males can distinguish females from other males, likely by detecting methyl-branched alkanes. These compounds are abundant in both males and newly-emerged females, rendering both unattractive to mating males. Third, methyl-branched alkanes are the only compounds that change in relative abundance in both females of chemotype 1 and chemotype 2. In contrast, alkenes differ between the two chemotypes, making the first hypothesis proposed in the introduction less likely, as males would require different receptors to perceive CHCs with double bonds at different carbon positions. Finally, the idea of methyl-branched hydrocarbons acting as anti-aphrodisiacs in very young females is more plausible, as we consistently recorded substantial quantities of unsaturated hydrocarbons in young females, making this compound less likely to trigger mating behavior in males.

Only specific methyl-branched alkanes seem likely to serve as anti-aphrodisiacs. The relative abundance of some methyl-branched alkanes decreases in both males and females (Fig. S2), making these compounds less likely candidates for this role. In contrast, 11-MeC23, 9-MeC23, 13-MeC25, 11-MeC25, and 9-MeC25 are promising candidates, as they represent the majority of the methyl-branched alkanes in both males and females. Notably, their relative abundance is high in males and in newly eclosed females, yet decreases in females, nearly disappearing three days after eclosion, while remaining high in males. Interestingly, these compounds are structurally similar, differing only by two carbons, suggesting they may be recognized by similar receptors and act as the primary methyl-branched alkanes with an anti-aphrodisiac function in *O. spinipes*. Changes in the abundance of methyl-branched alkanes in CHC profiles have been reported in other species, one of which has also been linked to female attractiveness.

Immature *Drosophila melanogaster* and *Drosophila simulans* express more methyl-branched alkanes than mature flies (Pechine et al. 1988). Female *Chrysomya putoria* flies (Braga et al. 2016) also express more methyl-branched and more dimethyl-branched alkanes, than mature flies. In *Nasonia vitripennis*, the abundance of methyl-branched alkanes in the female CHC profile correlates with female attractiveness. However, in contrast to *O. spinipes*, this correlation is positive (Sun et al. 2023).

The high abundance of methyl-branched alkanes in very young *O. spinipes* females could represent a mimicry of the male CHC profile, reducing male harassment of females before they are ready to mate. Mimicry of female CHC profiles by immature males to avoid harassment by other males has been reported in Hymenoptera and rove beetles (Peschke 1985, 1987b, a; Steiner et al. 2005). Avoidance of male harassment after mating has been shown in the digger wasp species *Stizus continuus* and is also associated with changes in female CHC profiles (Polidori et al. 2017).

In addition to a potential anti-aphrodisiac in eclosed females, a sex pheromone from 3-day-old females likely initiates the courtship behavior of the males. One candidate for the sex pheromone is the alkenes which increase significantly with age in females and are absent or present at lower levels in males. However, not the same ones increase in the two chemotypes: shorter alkenes (23-25 carbons) with double bond at uneven positions in chemotype 1 and longer ones (27-29 carbons) with double bonds at even positions in chemotype 2 (Fig. 4; Fig. S3; Moris et al. 2021; 2023). This makes it unlikely that a specific alkene is the pheromone, as has been shown for alkadienes with specific double bond positions and chain lengths in *D. melanogaster* (Ferveur 2005).

## Material and Methods

### Collection and rearing of Odynerus spinipes

We collected O. *spinipes* in 2016, 2018, and 2019 from trap nests installed in Büchelberg (Germany, Rhineland-Palatinate, WGS 84: 49.027985, 8.164801) (for more information, see (Moris et al. 2023)). We opened the trap nests in March and April to collect *O. spinipes* prepupae. The prepupae were stored in hard gelatin capsules (size 3, LUTOR, Cologne, Germany, and Birmingham, UK) at 5 °C for 2–4 weeks. Prior to the experiments, the gelatin capsules containing the prepupae were placed in a dark climate chamber at 23 °C to induce eclosion. To avoid sample loss due to desiccation, the gelatin capsules containing the prepupae were always kept in a closed polystyrene box with humidified peat, which we changed every two days. After hatching, the wasps were transferred to observation cages with a viewing window and zipper (30 cm x 30 cm x 30 cm; Bioform, Nuremberg, Germany), which we kept in a climate chamber at 21 °C, 70% relative humidity, and a 14.5/9.5 h day/night cycle. All wasps were fed with honey water in a petri dish lined with white paper. A total of 144 *O. spinipes* wasps were reared by us over three years. Of these, 40 were females with chemotype 1, 36 were females with chemotype 2, and 68 were males (most of them had their CHC profiles analyzed: see Table S4). In addition to wasps sampled in Germany, we collected female wasps in in 2017 in Tenneville (Belgium; WGS 84: 50.100671, 5.530061) and stored them in the freezer (−20°C) before further processing them so we could use them as dummies for behavioral experiments.

### Male mating behavior

To gain an understanding of the mating behavior of *O. spinipes* males, we recorded the behavior of nine *O. spinipes* males that had eclosed 4–5 days earlier, in the presence of age-controlled females. The females were kept separately in individual observation cages with food (paper with honey and water) in a climate chamber (70% humidity, 21 °C, 9.5 hours dark and 14.5 hours light) prior to the experiments. The males were kept under identical conditions, except that we kept them in pairs due to space constraints. The experiments were performed with wasps that had been exposed to 6.5 hours of light that day, which in mid-May in southern Germany corresponds to about 11:30 a.m., the time at which we observed samples of the species mating in the field. We started the experiments by placing a given female in a glass tube (inner diameter = 25 mm, height = 50 mm, volume = 25 ml) into which the male was introduced 5–10 s later. The tube was surrounded by white cardboards to create a uniformly illuminated environment for the wasps (Fig. 1A). A small hole in the cardboard on one side was used to record the wasp’s behavior using an EOS 5D Mark IV in video mode (Canon, Tokyo, Japan). Females and males were kept together and videotaped for 30 minutes. The behavior of the male wasp on the video recordings was then analyzed to derive an ethogram.

### Behavioral bioassays

To assess whether differential attraction to females of different ages is mediated by the CHC profile of the females, we conducted behavioral assays in which we offered *O. spinipes* males initially odorless dead female wasps that we coated with CHC extracts from 0-day-old and 3-day-old females. The females that we used as dummies were collected in 2017 in Tenneville (Belgium; WGS 84: 50.100671, 5.530061) and stored in the freezer (−20 °C) before further processing. The dummies were treated with dichloromethane in a Soxhlet extractor for at least ten cycles to remove their CHCs and other potential chemical cues. The wasps were then placed under a fume hood for 24 hours to ensure that the solvent had evaporated. To coat the dummies with the CHC profiles of females of a given age, we collected CHC extracts (see below for details) from 0-day-old and 3-day-old females reared in 2019 from the trap nests at the Büchelberg field site. An aliquot of each CHC extract was used for CHC profile characterization by GC-MS (see also Figure 3). The remaining volume of CHC extract was reduced to 200 µl by evaporation of excess solvent under a gentle nitrogen stream. The 200 µl of CHC extract was carefully applied drop by drop to the dummies over a time period of two days. The dummies were placed under a fume hood overnight to allow the solvent to evaporate. A total of 18 female dummies were prepared: seven were coated with CHC extract from 0-day-old chemotype 1 wasps, two with CHC extract from 0-day-old chemotype 2 wasps, seven with CHC extract from 3-day-old chemotype 1 wasps, and two with CHC extract from 3-day-old chemotype 2 wasps. The perfumed female dummies were presented to 4– 5-day old male wasps reared in 2019 from the same trap nests as the females whose CHC extracts we used to perfume the female dummies. To allow for the preparation of the dummies and the analysis of the CHC extracts used to coat them, the development of the males was delayed by keeping them at 5 °C for 1–2 weeks longer. Male wasps were kept isolated from females and from other males prior to the experiments.

Behavioral assays were performed as follows: a female dummy was placed with forceps (cleaned with dichloromethane between each experiment and dried under a fume hood) in the center of the bottom of an Erlenmeyer flask (50 mL volume, 85 mm height, 29 mm opening diameter) and surrounded by white cardboards (Fig. 1A). After 6.5 hours of light exposure on the day of the experiment, a male wasp was placed in the flask and its behavior was videotaped for 10 min., beginning with the first antenna contact of the male with the female dummy. After a break of 24 hours, we repeated the experiment, this time offering the male a female dummy coated with a different CHC extract. Half of the males were first presented with a dummy coated with a CHC extract from a 0-day-old female, followed by a dummy coated in a CHC extract from a 3-day-old female. For the other half, the order was reversed. We confirmed that the attractiveness scores did not differ significantly between dummies presented first and those presented second (Fig. S4). Each dummy was used only once. In order to have only the age of the female influence any differential attraction by the male, a given male was only provided with CHC extracts from females of the same chemotype. In total, we analyzed the behavior of nine males when they were presented sequentially with two dummies. In six of the nine trials, the male showed mating behavior. Only the behaviors from these trials were scored. Dummy attractiveness scores were calculated as two times the sum of the number of times a male exhibited behaviors B2–3 plus the sum of the number of times a male exhibited behavior B5. The weight of behaviors B2–3 was higher since these behaviors were only observed when the wasps mated while B5 was not always observed after mating and sometimes observed after unsuccessful mating attempts (Table 1). We compared the scores of attractiveness of the dummies using a paired t test performed in R (version 3.5.0 R Core Team, 2017).

### CHC profile analysis

We extracted the hydrocarbons and analyzed 149 CHC profiles of *O. spinipes* wasps of different age classes (1 [= 1d], 3 [= 3d], 8 [= 8d], 14 [= 14d] days after eclosion) and representing the two sexes and the two female chemotypes (male [= m], female chemotype 1 [= c1], female chemotype 2 [= c2]). Sample sizes were as follows: N_c1,0d_ = 9; N_c1,1d_ = 7; N_c1,3d_ = 7; N_c1,8d_ = 24; N_c1,14d_ = 8; N_c2,0d_ = 3; N_c2,1d_ = 9; N_c2,3d_ = 9; N_c2,8d_ = 24; N_c2,14d_ = 9; N_m,0d_ = 8; N_m,3d_ = 10; N_m,8d_ = 19; N_m,14d_ = 3: Table S4). In total we analyzed profiles from 40 females of chemotype 1, 36 females of chemotype 2, and 40 males since CHCs from female wasps collected in 2016 were sampled multiple times: 2–3, 7–8, and 10–14 days post eclosion using a solid-phase microextraction (SPME) fiber (Supelco, coating: polydimethylsiloxane, 100 µm; Sigma Aldrich, Bellefonte, PA, USA). One female of chemotype 1 was not sampled at 1-day-old because of time constraints due to multiple runs with the GCMS. To facilitate handling of the wasps sampled with SPME, they were anesthetized with CO_2_ and their metasoma was gently scrubbed with a SPME fiber for 2 minutes.

CHCs from wasps collected in 2018 and 2019 were sampled at day 0, day 3, and day 8 after eclosion. These wasps were killed by immersion in hexane for 2 min. in glass vials (Agilent Technologies, Santa Clara, CA, USA; screw-cap vials, 1.5 ml). The extracts were concentrated with a gentle stream of nitrogen to approximately 200 µL and then transferred to an insert vial (Agilent Technologies, Santa Clara, CA, USA; 250 ml glass with polymer feet).

CHC extracts on SPME fibers from wasps collected in 2016 were processed on an Agilent 6890 gas chromatograph coupled to an Agilent 5975 mass-selective detector (Agilent Technologies, Waldbronn, Germany). The GC was equipped with a DB-5 fused silica capillary column (30 m × 0.25 mm ID; df = 0.25 μm; J&W Scientific, Folsom, USA). The injector temperature was 250 °C. The following temperature profile was used: initial temperature of 60 °C, temperature increase of 20 °C per minute up to 150 °C, followed by a temperature increase of 5 °C per minute up to 300 °C, which was maintained for 10 min. CHC extracts in hexane solvent from wasps collected in 2018 and 2019 were analyzed using an Agilent 7890B gas chromatograph system (Agilent Technologies, Santa Clara, CA, USA) coupled to a 5977B mass-selective detector (Agilent Technologies, Santa Clara, CA, USA). The GC was equipped with a DB-5 column (30 m × 0.25 mm ID; df = 0.25 μm; J & W Scientific, Folsom, USA). The injector temperature was 250 °C. The following temperature profile was used: initial temperature of 40 °C, temperature increased of 10 °C per minute up to 300 °C, which was maintained for 10 minutes. Both systems used helium as the carrier gas at a constant flow rate of 1 mL/min. Electron ionization mass spectra (EI-MS) were acquired at an ionization voltage of 70 eV (source temperature: 230 °C). A standard solution of n-alkanes (C21–40) was injected weekly to control the sensitivity of the GC-MS and to provide a basis for the calculation of retention indices (RI).

Raw GC-MS data were analyzed using the MSD Enhanced ChemStation F.01.03.2357 software for Windows (Agilent Technologies, Böblingen, Germany). Individual compounds were characterized based on their diagnostic ions and their calculated Kovats retention index. The abundance of each CHC was normalized by dividing it by the total abundance of all CHCs. Identification of some methyl-branched alkanes was not possible because they were not properly separated on the above instruments with the above settings and were therefore treated as a mixture. The identified hydrocarbons and their average normalized abundance in the profiles of male and female wasps of different ages are listed in Table S2.

We only considered CHCs with a relative abundance of at least 0.3% in the CHC profile. To counteract the effects of normalization that created compositional data, we CLR-transformed the relative abundance data in the R package *compositions* (version 1.40-2) (Aitchison et al. 2000; van den Boogaart and Tolosana-Delgado 2008). We used non-metric multidimensional scaling (NMDS) and principal component analysis (PCA) to visualize the data in two dimensions. NMDS plot was performed using the 149 CHC profile analyzed to have a global view at the distribution of the different CHC profiles. Boxplots of the different CHCs were also carried using the 149 CHC profiles.

PCA was used to identify CHCs that differed most between 0-day-old and 3-day-old female wasps, regardless of their chemotype. For this purpose, we analyzed the CHC profile data of each chemotype separately and identified CHCs that occurred in at least one sample with a relative abundance of at least 1% and at the same time were highly correlated (≥ 0.7) in their variation across all samples in the PCA along the first principal component (axis 1). For these analyses, we used only CHC profiles of wasps sampled in 2019 to decrease the chance to pick up differences due to different GCMS or extraction ways (SPME, hexane extracts) rather than the ages.

NMDS plots were derived from Bray-Curtis distances between CHC profiles using the R packages *vegan* (version 2.4-2) (Oksanen 2011), *reshape2* (version 1.4.3) (Wickham 2007), and *dendextend* (version 1.5.2) (Galili 2015). PCAs were performed on the CLR-transformed abundance data using the R packages *FactoMineR* (Husson et al. 2017) and *Factoextra* (version 1.0.6) (Kassambara 2016).

Differences in the relative abundance of the identified CHCs between females of different age classes were assessed using a Wilcoxon rank correlation test and applying the Bonferroni correction to limit the number of false positives. The corresponding boxplots were drawn and statistical tests were performed in R (version 3.5.0; R Core Team, 2017) using the package *ggplot2* (version 3.2.1) (Wickham 2016).

## Acknowledgements

We thank Michael Schnell, Harald Noeske, and Rainer Blum for help with trap nest construction, Wolf Haberer for help with behavioral experiments, and Jean-Yves Baugnée for information on where to find *O. spinipes* in Belgium. We thank the Struktur-und Genehmigungsbehörde Süd and the Struktur-und Genehmigungsdirektion Nord (both Rhineland-Palatinate, Germany) for permission to collect samples and Gaby Schöning for permission to set up trap nests in a bee hotel in Büchelberg.

## Author contributions

O.N. and T.S. obtained funding for the project. V.C.M. conceived and designed the study. A.W. and V.C.M. conducted the behavioral studies. V.C.M. analyzed the behavioral data. V.C.M made the graphs and tables and figures. V.C.M. and T.S. performed the chemical analyses. O.N. and T.S. provided equipment and reagents. O.N. and V.C.M. wrote the main part of the manuscript. All authors provided critical feedback on the manuscript.

---

**Table S1:** Samples used and behaviors recorded in the behavioral bioassays with female dummies coated with CHC extracts from 0-day- and 3-day-old females. d = day(s).

| Male<br>CHC<br>ID | Male<br>ID | Female<br>CHC ID | Female<br>age | Chemotype | Female<br>presented | Mounting<br>attempt<br>(count) | Gaster<br>lifted<br>(count) | Aedeagus<br>extrusion<br>(count) | Attractiveness<br>score |
| --- | --- | --- | --- | --- | --- | --- | --- | --- | --- |
| 12942 | 1 | 12893 | 0 d | 1 | first | 1 | 0 | 0 | 2 |
| 12942 | 1 | 12951 | 3 d | 1 | second | 2 | 2 | 1 | 8 |
| 12944 | 2 | 12949 | 3 d | 1 | first | 2 | 1 | 0 | 5 |
| 12944 | 2 | 12891 | 0 d | 1 | second | 1 | 0 | 0 | 2 |
| 12940 | 3 | 12895 | 0 d | 1 | first | 1 | 0 | 0 | 2 |
| 12940 | 3 | 12953 | 3 d | 1 | second | 1 | 2 | 1 | 6 |
| 12952 | 4 | 12899 | 0 d | 1 | first | 3 | 0 | 0 | 6 |
| 12952 | 4 | 12969 | 3 d | 1 | second | 3 | 4 | 0 | 10 |
| 12954 | 5 | 12931 | 3 d | 2 | first | 3 | 0 | 0 | 6 |
| 12954 | 5 | 12909 | 0 d | 2 | second | 0 | 0 | 0 | 0 |
| 12958 | 6 | 12907 | 0 d | 1 | first | 2 | 1 | 2 | 9 |
| 12958 | 6 | 12975 | 3 d | 1 | second | 2 | 1 | 1 | 7 |

**Table S2:**
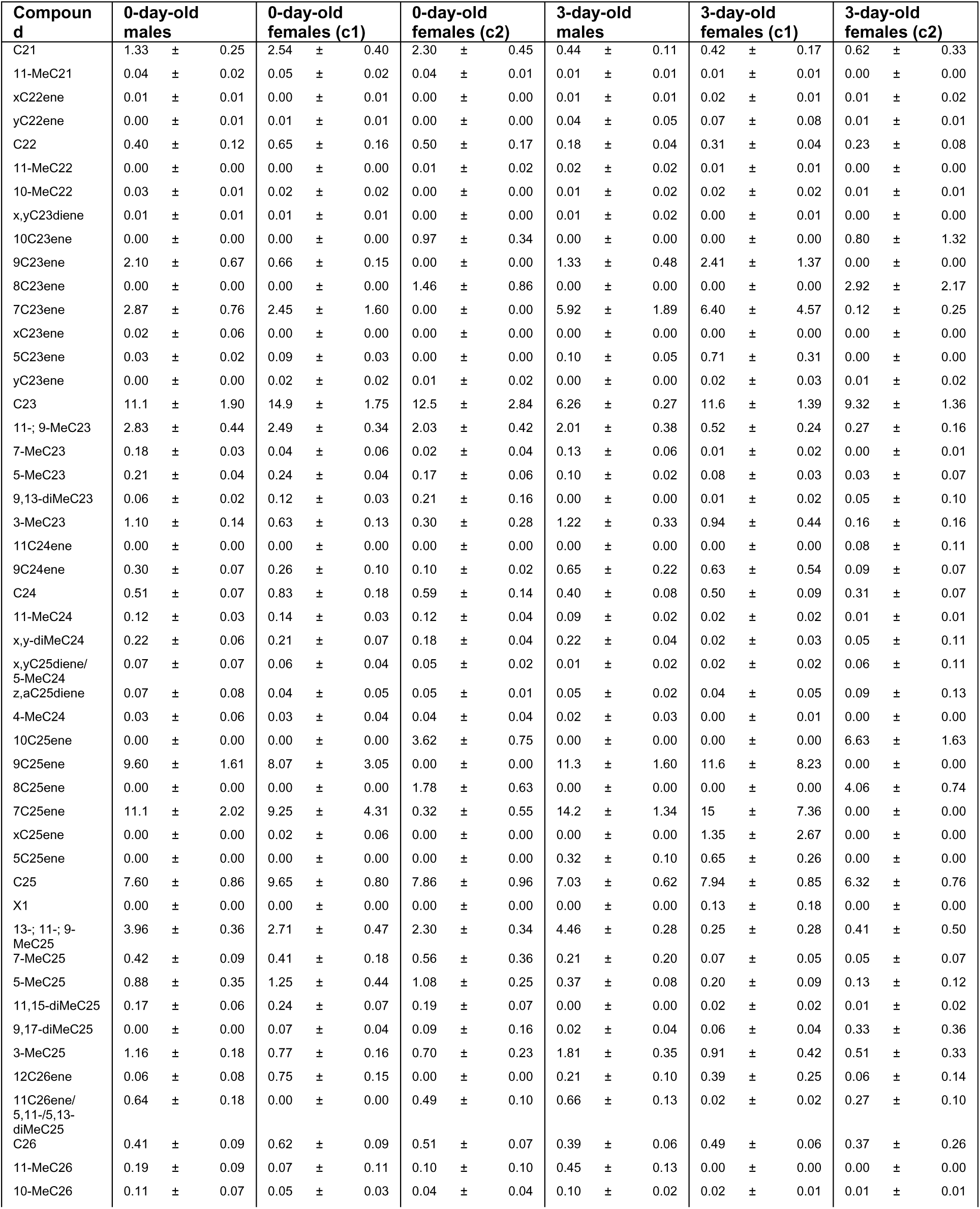
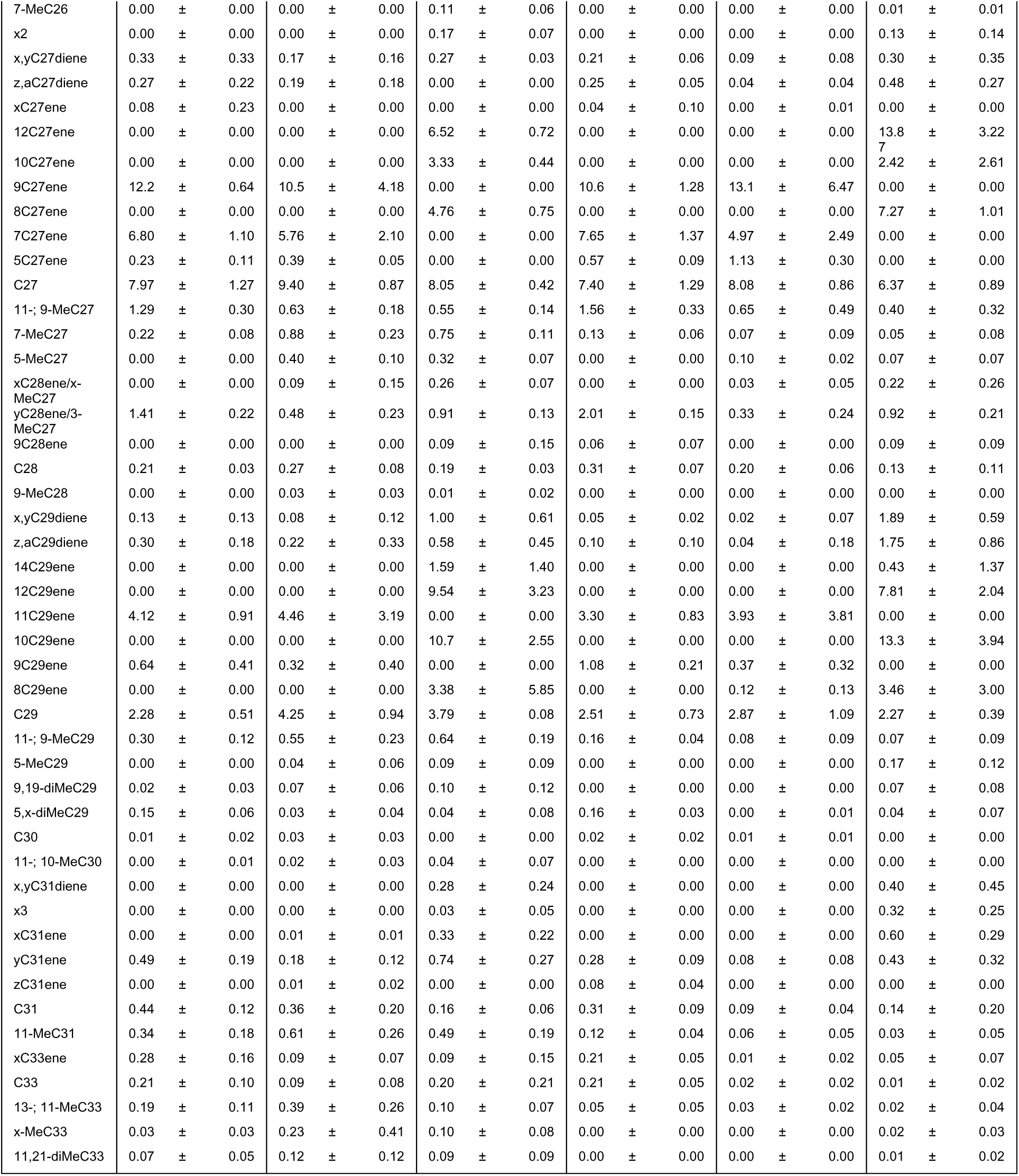
Relative abundances (% ± s.e.) of cuticular hydrocarbon compounds in extracts of *O. spinipes* males and females (c1 = chemotype 1; c2 = chemotype 2) on the day of eclosion and 3 days after eclosion.

**Table S3:** P-values of tests comparing the relative abundances of methyl-branched alkanes, which were found in PCA analyses to decrease in females of chemotype 1 (= c1) and chemotype 2 (= c2), but not in males. P-values were Bonferroni-corrected for multiple testing within each group.

|  | Females c1 |  | Females c2 |  | Males |  |
| --- | --- | --- | --- | --- | --- | --- |
|  | p | adjusted p | p | adjusted p | p | adjusted p |
| 11-; 9-MeC23 | 0.0001748 | 0.000874 | 0.009091 | 0.045455 | 0.0008684 | 0.004342 |
| 7-MeC27 | 0.000998 | 0.00499 | 0.01441 | 0.07205 | 0.01404 | 0.0702 |
| 13-; 11-; 9-MeC25 | 0.0009471 | 0.0047355 | 0.009091 | 0.045455 | 0.4082 | >1 |
| x,y-diMec24 | 0.000998 | 0.00499 | 0.05981 | 0.29905 | 0.2743 | >1 |
| sum of methyl branched | 0.0001748 | 0.000874 | 0.009091 | 0.045455 | 0.2031 | >1 |

**Table S4:** Number of CHC profiles analyzed for *O. spinipes* females of chemotype1 (= c1), of chemotype 2 (= c2), and males at specific ages (0 days, 1 day, 3 days, 8 days, and 14 days old), sampled in 2016, 2018, and 2019 from trap nests collected in Büchelberg.

|  | 2016 | 2018 | 2019 | Total |
| --- | --- | --- | --- | --- |
| **Females C1** |  |  |  | **55** |
| 0d |  |  | 9 | 9 |
| 1d | 7* |  |  | 7 |
| 3d |  |  | 7 | 7 |
| 8d | 8* |  | 16 | 24 |
| 14d | 8* |  |  | 8 |
| **Females C2** |  |  |  | **54** |
| 0d |  |  | 3 | 3 |
| 1d | 9* |  |  | 9 |
| 3d |  | 5 | 4 | 9 |
| 8d | 9* | 10 | 5 | 24 |
| 14d | 9* |  |  | 9 |
| **Males** |  |  |  | **40** |
| 0d |  |  | 8 | 8 |
| 3d |  | 3 | 7 | 10 |
| 8d |  | 3 | 16 | 19 |
| 14d |  | 3 |  | 3 |
| **Total** | **50** | **24** | **75** | **149** |

\* Samples observed throughout their lifespan, with some having their hydrocarbons sampled three times. Seven females of chemotype 1 and nine females of chemotype 2 had their CHC profiles sampled three times. One female of chemotype 1 was sampled twice.

**Fig. S1:**
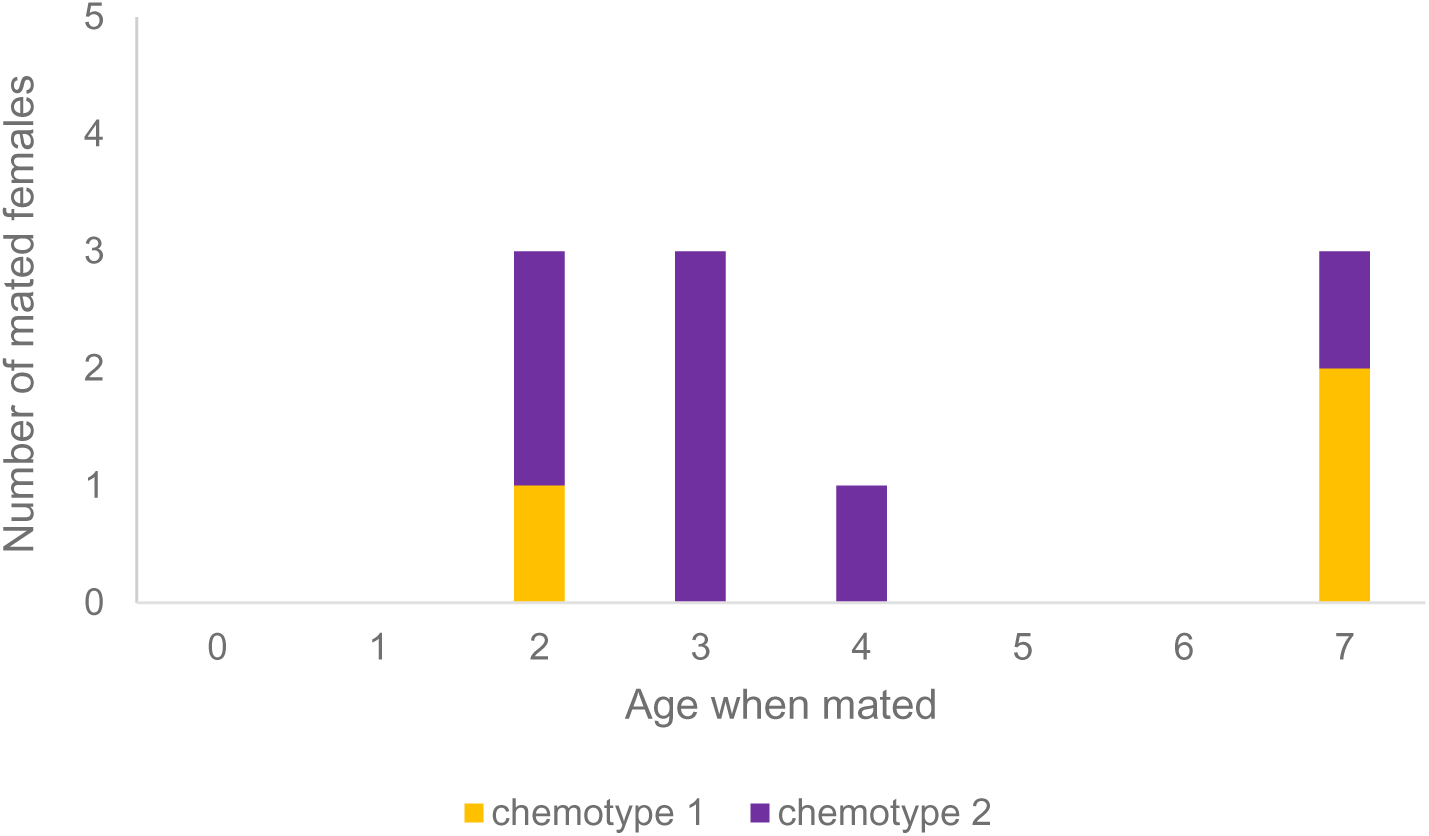
Number of *O. spinipes* females of chemotype 1 (orange) and chemotype 2 (violet) observed to be mated at specific ages (x-axis).

**Fig. S2:**
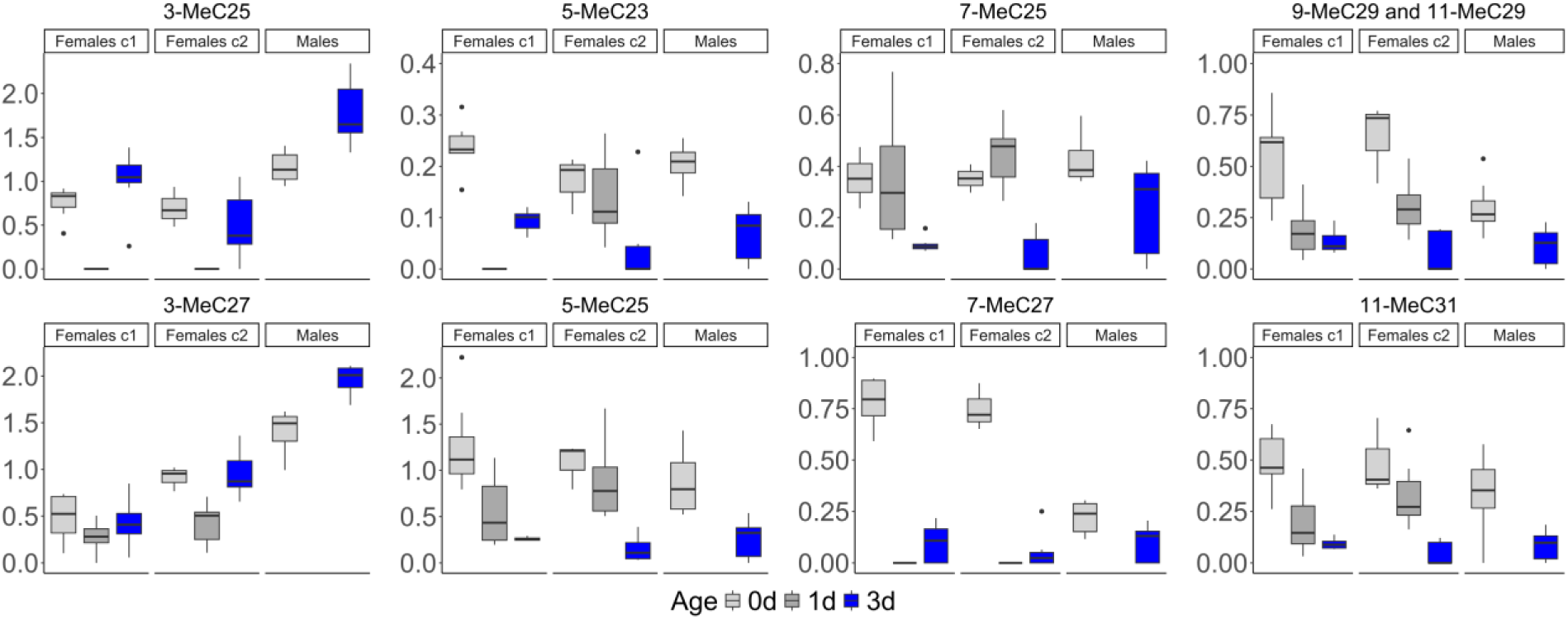
Relative abundance of methyl-branched alkanes in *O. spinipes* males and females (c1 = chemotype 1; c2 = chemotype 2) on the day of eclosion (gray) and 3 days after eclosion (blue). The ordinate shows the relative abundance (in percent) of the different methyl-branched alkanes.

**Fig. S3:**
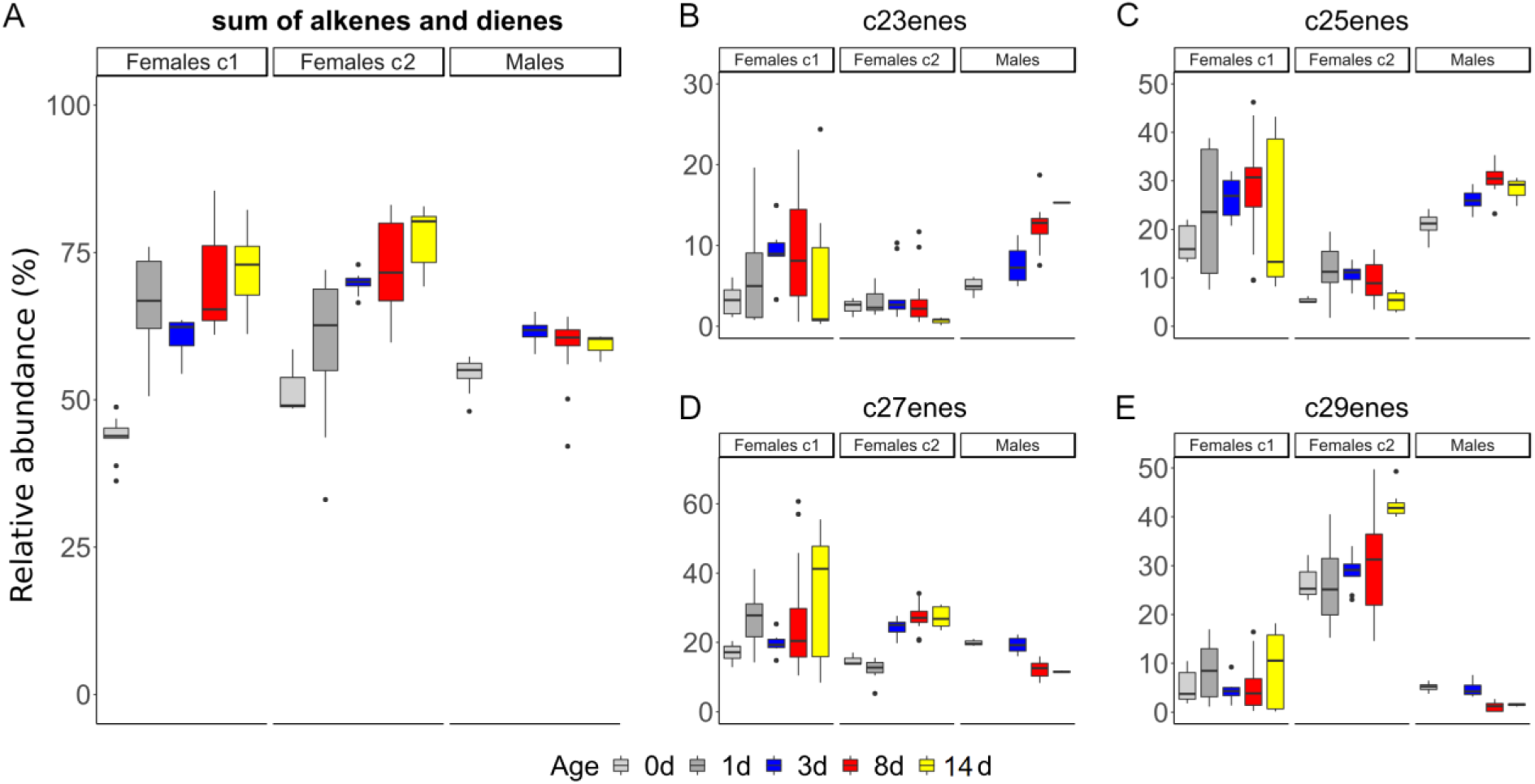
Relative abundance of alkenes and alkadienes in *O. spinipes* females of chemotype 1 (Females c1); in *O. spinipes* females of chemotype 2 (Females c2) and in *O. spinipes* males (Males) at day 0 (grey), day 1 (dark grey), day 3 (bleu), day 8 (red), and day 14 (yellow). y axis is the relative abundance (in percent) of all alkenes and dienes (A), of all alkenes and dienes with a chain length of 23 carbons (B), of all alkenes and dienes with a chain length of 25 carbons (C), of all alkenes and dienes with a chain length of 27 carbons (D), of all alkenes and dienes with a chain length of 29 carbons (E).

**Fig. S4:**
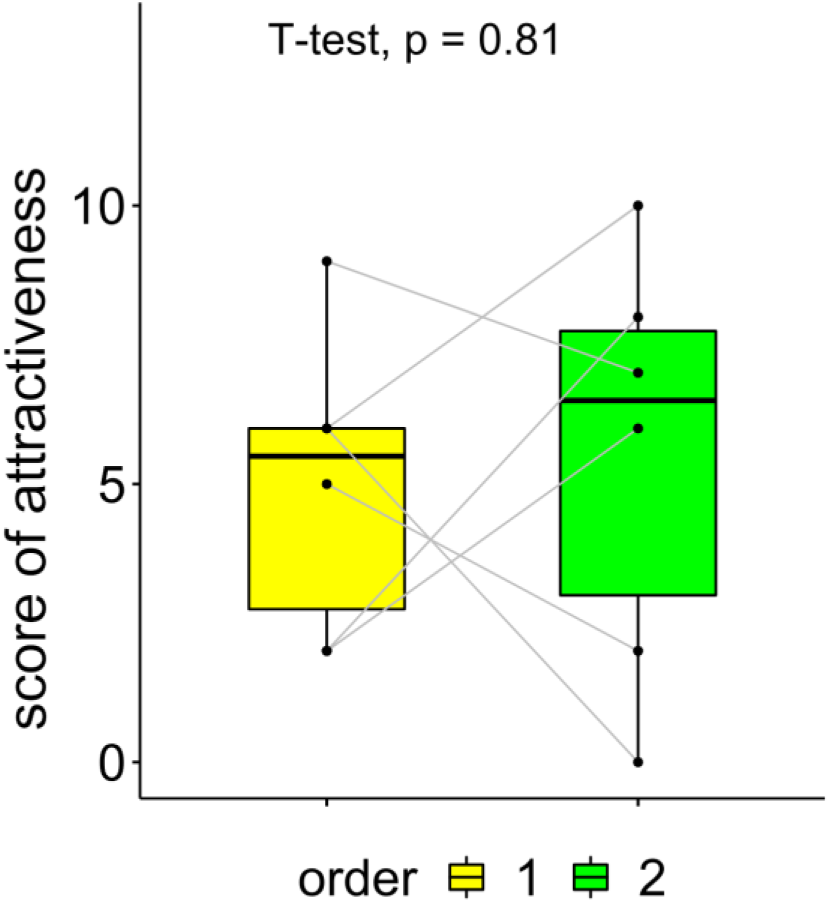
Attractiveness scores of female dummies coated with cuticular hydrocarbons extracted from females presented to six males, as first dummy (yellow) and as second dummy (green). Gray lines indicate each pair of data. See Materials and methods for how attractiveness scores were calculated.

